# Inferring Dynamic Information from Protein Structures by Gaussian Integrals and Deep Learning

**DOI:** 10.1101/2025.09.22.677755

**Authors:** Felipe Vilicich, Zhaoqian Su, Shanye Yin, Yinghao Wu

## Abstract

Protein conformational flexibility underlies a wide range of biological functions, yet experimentally probing dynamics at atomic resolution remains costly and low-throughput. Here, we present a deep learning framework that predicts protein flexibility directly from static structural descriptors, bypassing the need for molecular dynamics (MD) simulations. Using the ATLAS database of standardized all-atom MD trajectories, we encoded 1,374 protein chains as 30-dimensional Gaussian integral (GI) vectors—global shape and topology invariants of the protein backbone. Principal component analysis of GI profiles revealed four structural clusters with distinct secondary structure compositions and flexibility distributions. We trained an attention-based one-dimensional convolutional neural network (1D-CNN) to classify proteins as flexible or non-flexible based on their root-mean-square fluctuation (RMSF) relative to the dataset-wide mean. The classifier achieved an AUC of 0.772 (95% CI: 0.712–0.826) on an independent test set, with balanced sensitivity and specificity, and identified a small subset of GI components as the most predictive. In a regression setting, a recurrent neural network outperformed other architectures, attaining an R^2^ of 0.537, though high-flexibility values were systematically underestimated. Cluster-specific analyses indicated that coil-rich and β-sheet–dominated proteins were more amenable to flexibility prediction than α-helical proteins, likely due to greater structural heterogeneity. Our results demonstrate that compact GI descriptors preserve sufficient information to recover MD-derived flexibility trends, offering a computationally efficient complement to simulation-based approaches. This framework enables large-scale screening of protein dynamics from structural data alone, with potential applications in structural bioinformatics, drug design, and functional annotation.

## INTRODUCTION

Proteins are essential components in most biological processes in living organisms^1^. The majority of these biomolecules adopt specific three-dimensional structures, which are determined by their primary sequences^2^. However, protein structures are not static under physiological conditions and undergo continuous conformational changes^3^. This dynamic behavior allows proteins to carry out their functions in the cellular environment^4^. Conformational variations enable proteins to switch between different functional states, allowing them to respond dynamically to external signals and enhancing their affinities for substrates or interactions with other molecules. These capabilities are fundamental to processes such as enzyme regulation and signal transduction. Therefore, exploring protein dynamics and how they are influenced by protein structure is crucial for gaining deeper insights into their roles in biology.

Information on protein dynamics can be characterized experimentally by methods such as nuclear magnetic resonance (NMR)^5^ and fluorescence resonance energy transfer (FRET)^6^. Nevertheless, the spatial-temporal resolutions of the dynamic information obtained by these methods are limited, preventing them from providing mechanistic insights of protein functions at atomic detail. Moreover, these experimental approaches are time-consuming and labor-intensive, making them unsuitable for large-scale studies or high-throughput screening. Finally, different experimental techniques require proteins to be studied under varying conditions, which may not accurately replicate the physiological environment in which these proteins function in living organisms. This discrepancy poses a challenge in interpreting experimental results and comparing outputs across different techniques.

Computational methods serve as an effective alternative for testing conditions that are currently inaccessible in the laboratory. Among a large variety of computational tools available, molecular dynamics (MD) simulation is specifically designed to study the physical movements and interactions of biomolecules over time^7-14^. By solving Newton’s equations of motion for each particle in a system, this method simulates molecular dynamics at the atomic level. MD simulations have been widely applied to evaluate the effects of conformational changes on enzyme catalysis, allosteric pathways, ligand-receptor recognition, and protein complex assembly^15^. Despite their versatility, MD simulations require significant computational resources. Systematic testing to simulate large numbers of biomolecules over long time scales is impractical without access to a limited number of high-performance computing facilities, such as Anton^16^. Furthermore, variations in software, force fields, system settings, or simulation protocols can produce inconsistent results^17^, even for the same proteins, making it extremely difficult to compare results from different research groups. As a result, there is a growing demand for computationally efficient tools that can systematically analyze protein conformational dynamics. Such tools should complement MD simulations and provide significant value to the scientific community by advancing the understanding of protein function.

Machine learning could be an ideal option for carrying out this task. It is a highly developing branch of artificial intelligence in which a computational model is trained to identify patterns and make predictions from specific datasets^18^. Given the growing amount of data available in the protein data bank (PDB)^19^, the application of machine learning, especially deep learning algorithms, to structural biology has achieved significant success^20^. Methods such as AlphaFold^21,22^ and RoseTTAFold^23^ achieve high accuracy in predicting protein tertiary structures. AlphaFold uses multiple layers of attention mechanisms to predict the distance between pairs of amino acids and the angles of their chemical bonds by learning from a vast dataset of known protein structures and their sequences. These methods have recently been extended to model the structures of multimeric protein complexes. In contrast, analyzing and predicting protein dynamics using machine learning remains a major challenge, primarily due to the scarcity of reliable data.

In this paper, we present a deep learning framework to infer dynamic information from protein structures. By leveraging a recently developed database, ATLAS^24^, we can access the all-atom MD simulation results for more than one thousand proteins with diverse ternary structures. We further transferred these protein structures into high-dimensional vectors by Gaussian integrals (GIs)^25^. The vectors were fed to a 1D convolutional network with an attention layer to predict the amplitudes of protein conformational fluctuations. Based on the cross-validation against the ATLAS database, we showed that our deep learning model can predict whether the flexibility of a given protein is higher or lower than a predefined threshold with more than 70% accuracy. Using a regression layer as the output, we also reproduced the specific values of root-mean-square-fluctuations (RMSF)^26^, which formed a strong correlation with the MD simulation data. Because GIs are descriptors of protein shape, our study demonstrated that the global dynamics of a protein can be decoded by deep learning without knowing its sequence information. This suggests that conformational flexibilities of proteins are embedded in their structures. In summary, the development of this computationally efficient tool can compensate for the current limitation of MD simulations and serve as a useful addition to a suite of experimental approaches that study protein dynamics.

## METHODS

### Dataset

All data used in this study were drawn from the ATLAS database of standardized all-atom molecular dynamics (MD) simulations^24^. ATLAS comprises trajectories for 1,390 protein chains representing 1,149 non-redundant ECOD X-class domains, each simulated in triplicate for 100 ns using GROMACS with the CHARMM36m force field. From these trajectories, per-residue RMSF of Cα atoms were computed and averaged across replicates to yield a single RMSF value per protein, which serves as our target variable (**Fig. 1A**).

**Figure 1.**
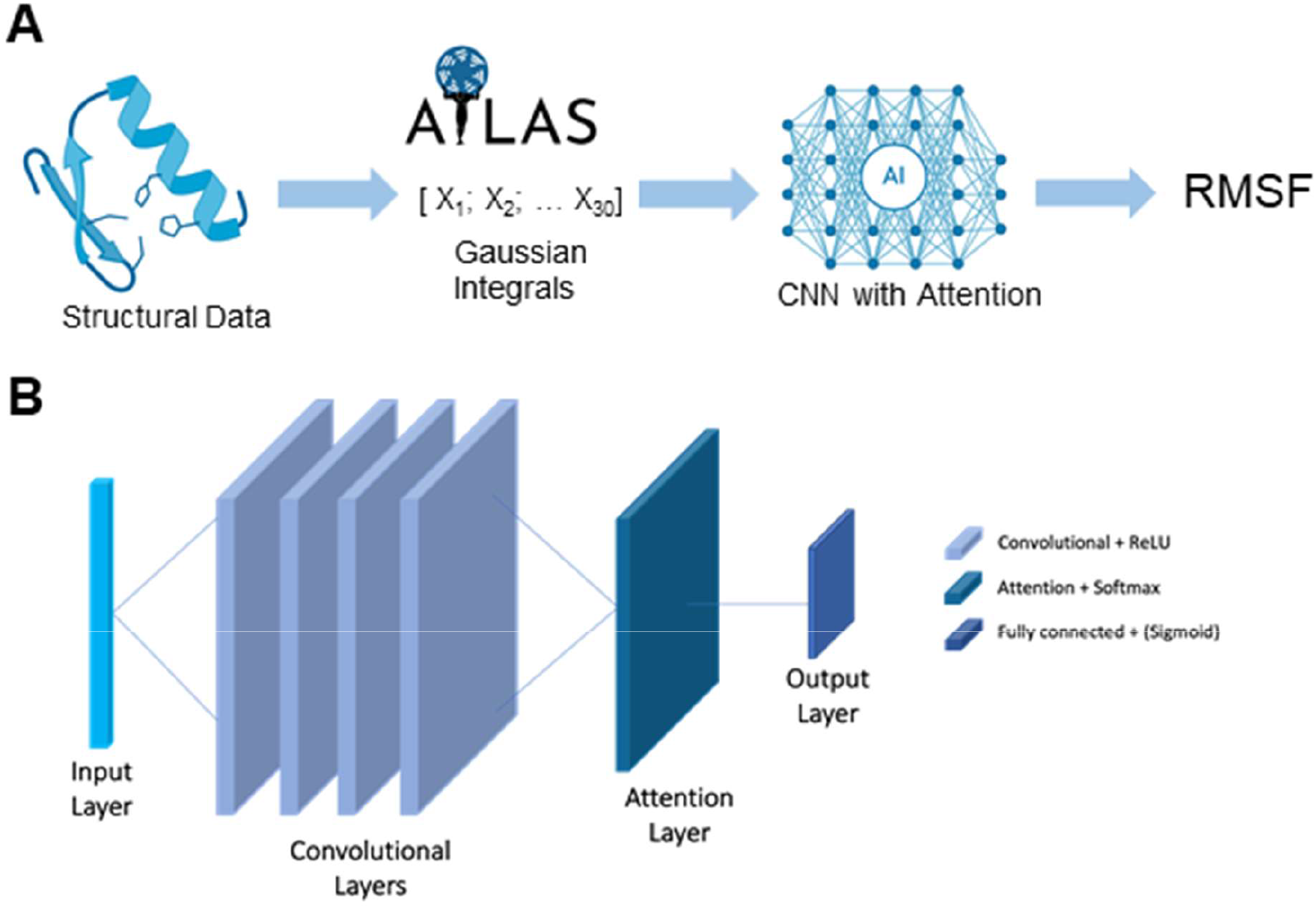
Outline of the problem. **A)** Flow chart showing our approach to solve the problem. **B)** Architecture of the model for regression and classification tasks.

### Gaussian integrals

To transform 3D structures into fixed-length feature vectors, we computed 30 topological invariants, also known as generalized Gauss integrals, capturing global curve properties of the protein backbone (writhe, average crossing number, and higher-order correlations)^25^. These Gauss integral descriptors, normalized to unit variance across the dataset, serve as input features to our convolutional network with attention (**Fig. 1A**).

### Deep learning models

Our classifier is a one-dimensional convolutional neural network (1D-CNN) with an embedded attention mechanism^27^ applied to the Gaussian integral (GI) sequence. Specifically, the model treats each GI vector as a single-channel time series of length 30, resulting in input tensors of shape (batch size × 1 × 30). The network consists of four stacked 1D convolutional blocks (kernel size = 3, padding = 1, no dilation), each followed by a ReLU activation, enabling the learning of hierarchical patterns across the sequence. An attention layer is then applied to compute learned importance weights over the 30 positions, aggregating them into a single 128-dimensional feature vector. This vector is passed through a fully connected layer that maps it to a single logit, which is then transformed via a sigmoid activation to produce a probability score (**Fig. 1B**). The attention mechanism was included to allow the model to learn which positions along the GI sequence are most informative for classification. Unlike traditional pooling methods that treat all positions equally, attention dynamically weights each of the 30 input positions based on their contribution to the final prediction. This not only improves performance by focusing the model’s capacity on relevant features, but also enhances interpretability by enabling us to probe which regions of the structural encoding influence the classifier’s decision.

### Training and validation

Five-fold cross-validation was performed on the 1,099 proteins (with a randomly held-out test set of 275). In each fold, 80% of the 1,099 served for training and 20% for validation, stratified by class (flexible and non-flexible). We used BCELoss, Adam (learning rate = 10^−3^, weight_decay = 10^−4^), batch size = 32, and early stopping (patience = 7, max epochs = 50). After identifying the fold with the best validation accuracy, we retrained that model on all 1,099 examples and evaluated its final performance on the independent 275-protein test set. Loss curves for training and validation can be found in supplementary section.

### Data and code availability

All data from this study, including the can be found in the GitHub repository https://github.com/fvilicich/gaussian_integral/blob/main/gaussian_integral_classification_vfinal.ipynb.

## RESULTS

### Statistical description of ATLAS database

We first performed a principal component analysis (PCA)^28^ of the GIs calculated for 1374 protein chains from the ATLAS database, revealing four distinct clusters, labeled 1 through 4. The first two principal components (PC1 and PC2) capture 29.53% and 20.26% of the total variance in the dataset, respectively, indicating that nearly half of the structural variability encoded by the GIs can be visualized in this two-dimensional projection (**Fig. 2A**).

**Figure 2.**
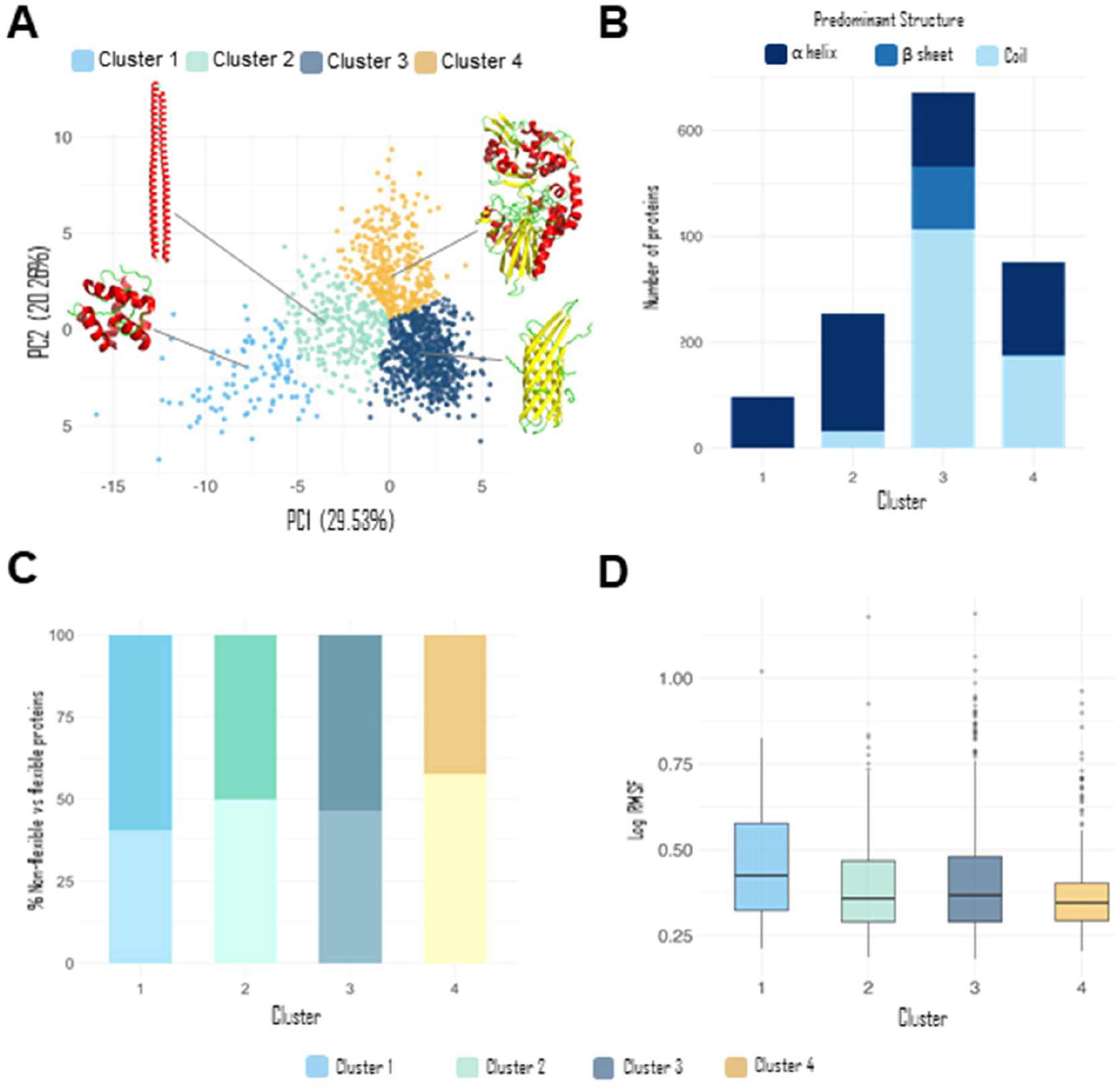
Statistical description of ATLAS database. **A)** Principal component analysis of the Gaussian integral vectors with schematic protein structures. **B)** Bar plot illustrating the predominant secondary structures on each cluster in the PCA space. **C)** Bar plot illustrating the proportion of non-flexible and flexible proteins on each cluster in the PCA space. Darker shades of color represent flexible proteins and lighter shades represent non-flexible proteins. **D)** Boxplot of the RMSF values for each cluster in the PCA space.

Then, we examined the structural composition of each cluster in terms of secondary structure content. The annotations for α-helix, β-sheet, and coil were obtained directly from the ATLAS database, which provides residue-level assignments for each protein chain. To assign a single predominant structure to each protein, we applied a simple majority rule: the secondary structure type with the highest residue count along the chain was considered the dominant one. For example, a protein composed of 70% α-helix, 20% β-sheet, and 10% coil was categorized as α-helical. Such classification scheme, while reductive, enables a tractable comparison of structural trends across clusters and reveals distinct secondary structure preferences associated with each group. This analysis showed that clusters 1 and 2 are enriched in α-helical proteins, while cluster 3 is dominated by coil-rich proteins, with additional representation from β-sheets and α-helices. Cluster 4 displays a near-even split between α-helical and coil-dominant proteins (**Fig. 2B**). These patterns support the idea that the GI representation captures structural features relevant to secondary structure content and provide a foundation for exploring whether such encodings can predict functional characteristics.

We next assessed the distribution of protein flexibility across clusters by classifying proteins as either flexible or non-flexible, based on their RMSF derived from ATLAS. Proteins with RMSF values above the dataset-wide mean were labeled as flexible, whereas those with values below the mean were considered non-flexible. Clusters 2 and 3 exhibited a nearly balanced distribution of flexible and non-flexible proteins. Cluster 1 showed a slight enrichment of flexible proteins (∼57%), whereas Cluster 4 was predominantly composed of non-flexible proteins (**Fig. 2C**). Notably, the overall dataset displayed an approximately even split between flexible and non-flexible proteins, serving as a useful baseline for interpreting cluster-specific deviations. To further investigate these differences, we plotted the log-transformed RMSF values per cluster (**Fig. 2D**). Cluster 1, which contains the highest proportion of flexible proteins (as shown in **Fig. 2C**), also exhibits the highest median RMSF, supporting its characterization as structurally more dynamic. In contrast, Cluster 4, which is predominantly composed of non-flexible proteins, shows the lowest median RMSF, suggesting a more rigid structural profile.

### Predicting protein flexibility

We next assessed whether GI descriptors could be used to predict protein flexibility, framing the task as a binary classification based on their RMSF values (above or below the dataset-wide mean, as in **Fig.2C**). We held out a test set of 275 randomly selected proteins and used the remaining 1,099 proteins to train a 1D convolutional network with an embedded attention mechanism by 5-fold cross-validation. The receiver operating characteristic (ROC)^29^ curve (**Fig. 3A**) yielded an AUC of 0.772 (95% CI: 0.712–0.826), indicating good discriminative performance on the test set. The optimal classification threshold, determined by the Youden J statistic^30^, achieved a true positive rate of ∼0.67 and a false positive rate of ∼0.23. The precision–recall (PR) curve (**Fig. 3B**) resulted in an average precision of 0.797, with a maximum F1 score of 0.726 at a recall of ∼0.63 and precision of ∼0.72, suggesting a favorable balance between sensitivity and specificity. The confusion matrix (**Fig. 3C**) indicated that the classifier correctly identified 75.0% of non-flexible proteins and 69.1% of flexible proteins. Calibration analysis (**Fig. 3D**) showed moderate alignment between predicted probabilities and observed frequencies, with slight underestimation in the lower probability bins.

**Figure 3.**
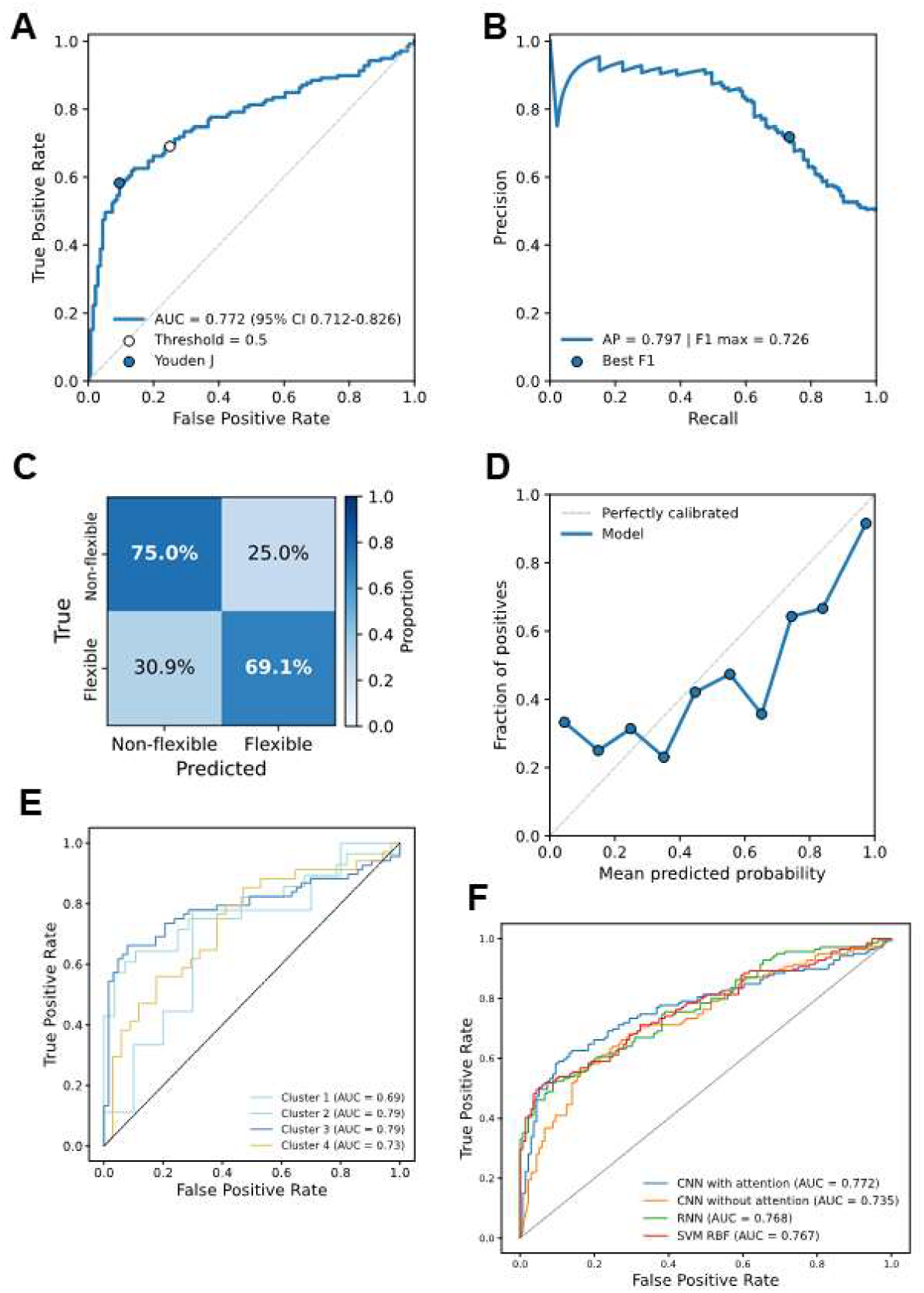
Predicting protein flexibility from GI descriptors using deep learning. **A)** Receiver operating characteristic (ROC) curve of the CNN with attention on the independent test set (n = 275). The optimal threshold (Youden J statistic) is indicated. **B)** Precision–recall (PR) curve of the same model. **C)** Normalized confusion matrix showing classification performance. **D)** Calibration curve comparing predicted probabilities to observed frequencies. **E)** Cluster-specific ROC curves demonstrating performance across structural groups. **F)** Model comparison of the CNN with attention against alternative architectures.

Cluster-specific ROC curves (**Fig. 3E**) revealed that predictive performance was highest for clusters 2 and 3 (AUC = 0.79 for both), followed by cluster 4 (AUC = 0.73) and cluster 1 (AUC = 0.69). As noted in **Fig. 2B**, clusters 2 and 3 contain higher proportions of coil-rich and β-sheet proteins, respectively, which may be more amenable to flexibility prediction. In contrast, the poorer performance for cluster 1 could be influenced by its strong enrichment in α-helical proteins and its small sample size in the testing set (n = 19), both of which may limit predictive resolution.

We also compared the CNN with attention to three alternative models: a CNN without attention, a recurrent neural network (RNN)^31^, and a support vector machine with radial basis function kernel (SVM RBF)^32^. The CNN with attention achieved the highest AUC (0.772), outperforming the CNN without attention (0.735), RNN (0.768), and SVM RBF (0.767) (**Fig. 3F**), supporting the benefit of incorporating attention mechanisms for this task.

As additional analysis, we further compared the distributions of predicted probability for proteins labeled as non-flexible (class 0) and flexible (class 1) across four structural clusters. As shown in **Supplementary Fig. 1A**, we found that predictions for non-flexible (Class-0) proteins are consistently more compact than for flexible (Class 1) proteins, as indicated by the smaller interquartile ranges across clusters. The broader Class 1 spread likely reflects heterogeneous structural mechanisms of flexibility.

Finally, training and validation loss curves across epochs are shown in **Supplementary Fig. 1B**. The CNN with attention exhibited progressive reduction in binary cross-entropy loss for both sets, although validation loss plateaued after ∼20 epochs, with a late increase indicating potential overfitting. Feature attribution analysis using Integrated Gradients (**Supplementary Fig. 1C**) revealed that only a small subset of GI components—particularly positions 1, 2, 3, 4, 7, 11, and 15—contributed disproportionately to classification, suggesting that specific regions of the GI descriptor encode most of the flexibility-related signal.

### Regression analysis of RMSF values

In addition to binary classification, we evaluated the predictive capacity of the GI descriptor in a regression setting, training the model to estimate the continuous RMSF value of each protein. Across the test set, predicted and true RMSF values displayed a moderately strong correlation coefficient of 0.68 (R^2^ = 0.46, slope = 0.70; **Fig. 4A**). While the model captured the overall trend, predictions tended to underestimate higher RMSF values, indicating reduced sensitivity for highly flexible proteins.

**Figure 4.**
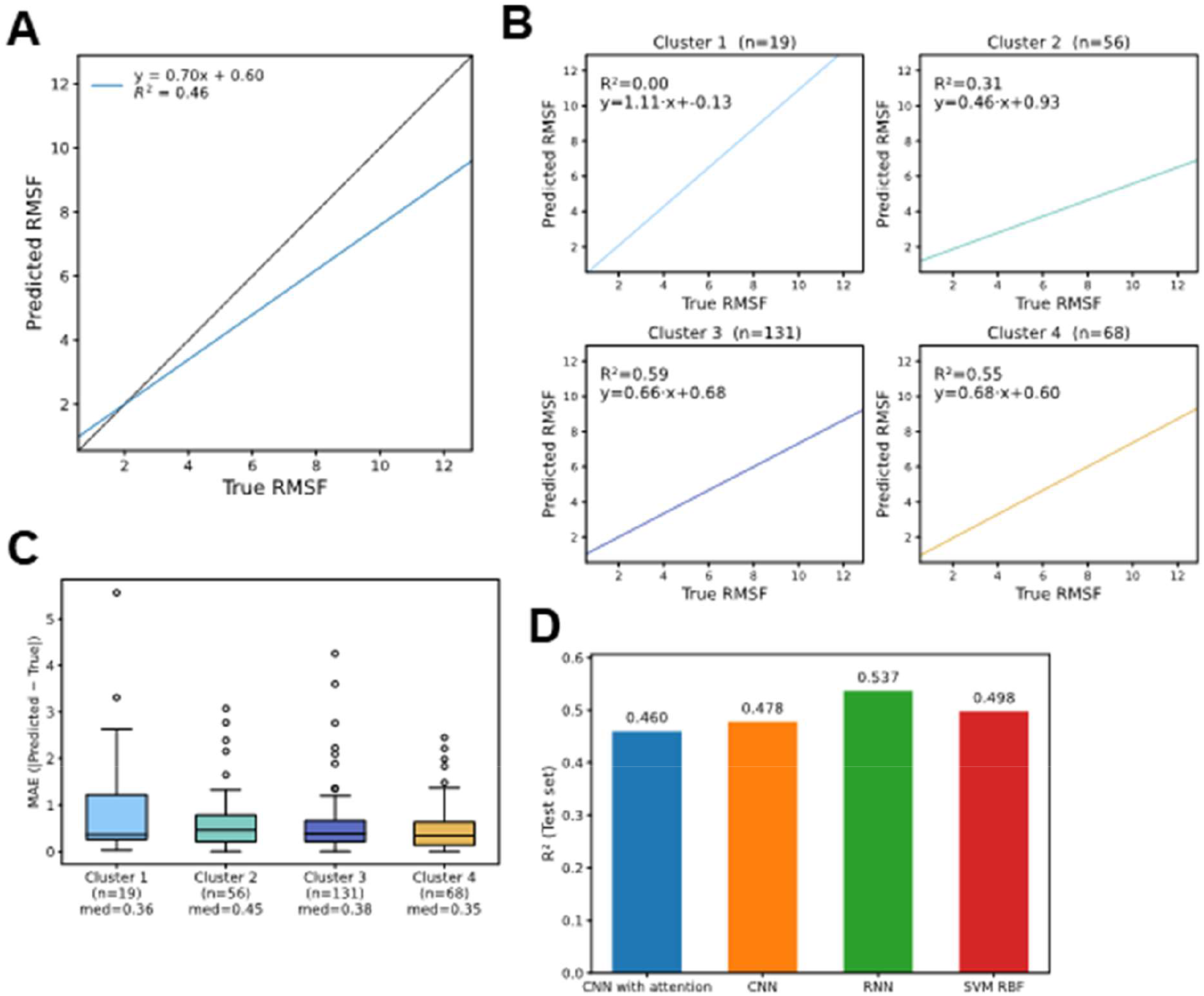
Regression analysis of protein flexibility from GI descriptors. **A)** Scatter plot of predicted versus true RMSF values with linear regression fit. **B)** Cluster-specific scatter plots showing predicted versus true RMSF for each structural group. **C)** Box plots of median absolute error (MAE) across clusters. **D)** Comparison of regression performance (R^2^ on test set) across different model architectures.

The distribution of predominant secondary structures across clusters (**Fig. 2B**) may partly explain the variability in predictive performance observed in the regression analysis (**Fig. 4B**). Clusters 3 and 4, which exhibited the highest R^2^ values in the regression task, contain substantial proportions of coil-rich and β-sheet proteins, respectively. These secondary structure types may encode GI patterns that more strongly correlate with RMSF, facilitating better prediction. In contrast, cluster 1—predominantly α-helical—showed no measurable correlation between predicted and true RMSF values, and also contained the fewest proteins, compounding the difficulty of establishing a robust mapping. Cluster 2, also enriched in α-helical proteins, achieved only moderate performance, suggesting that the more repetitive and rigid nature of α-helical folds could limit the variability captured by GIs for flexibility prediction.

Notably, cluster 1 contained only 19 proteins, and cluster 2 contained 56 proteins, which likely limits the statistical robustness of these correlation estimates compared to clusters 3 (n = 131) and 4 (n = 68). This imbalance in sample size should be considered when interpreting cluster-specific regression metrics. Median absolute error (MAE) values ranged from 0.35 to 0.45 across clusters, with cluster 4 showing the lowest median error (0.35) and cluster 2 the highest (0.45) (**Fig. 4C**). Although the error distribution was relatively consistent, a subset of proteins in each cluster showed large deviations, proving the difficulty of accurately predicting extreme RMSF values.

Model-to-model comparison (**Fig. 4D**) revealed a reversal in performance trends compared to the classification task (**Fig. 3F**). While the CNN with attention was the top performer for binary classification, the regression task was best handled by the recurrent neural network (RNN), which achieved the highest R^2^ on the test set (0.537). This suggests that regression benefits more from the RNN’s ability to capture sequential dependencies across the GI descriptor, whereas classification gains more from the attention mechanism’s capacity to focus on a small subset of highly informative positions.

Training dynamics are shown in **Supplementary Fig. 2B**. The model exhibited progressive loss reduction for both training and validation sets, with validation loss plateauing after ∼20 epochs before a late increase, suggesting mild overfitting. The relationship between predicted and true RMSF values for proteins with RMSF ≤ 5 (**Supplementary Fig. 2A**) yielded the same R^2^ value as the full dataset (0.46), but variance was lower in the low-flexibility range.

## CONCLUDING DISCUSSIONS

This study demonstrates that Gaussian integral (GI) descriptors capture meaningful structural variability across a large and diverse protein set, and that these representations can be leveraged to predict protein flexibility in both classification and regression contexts. The principal component analysis of GI profiles revealed four distinct structural clusters, which differed not only in predominant secondary structure composition but also in flexibility distribution. Notably, clusters enriched in α-helical proteins (clusters 1 and 2) were generally less predictable in terms of flexibility, while coil-rich (cluster 3) and β-sheet–dominated (cluster 4) groups exhibited more consistent predictive performance. These trends suggest that the repetitive geometry and rigidity of α-helices may limit the variability encoded by GIs, whereas the more heterogeneous topologies of coils and β-sheets produce richer flexibility signatures.

One of the most striking aspects of these findings is that they are derived from an extremely compact representation: a 30-element GI vector per protein. Despite their simplicity, these descriptors—originally developed as global shape and topology encodings—retain sufficient resolution to recover trends typically extracted from computationally intensive molecular dynamics (MD) simulations. In essence, we are able to approximate dynamic information such as relative flexibility (RMSF) directly from static structural descriptors, bypassing the need for atomistic time evolution while preserving predictive relevance. This underscores the potential of low-dimensional, information-rich embeddings for biophysical property prediction and opens opportunities for large-scale flexibility screening across proteomes without the prohibitive cost of full MD trajectories.

In the binary classification task, the attention-based CNN achieved the best performance, with an overall AUC of 0.772 and balanced sensitivity–specificity trade-offs. The attention mechanism likely enabled the model to identify a small subset of GI positions that were disproportionately informative for discriminating flexible from non-flexible proteins, as confirmed by feature attribution analysis. This is consistent with the observation that specific structural features— potentially reflecting localized mobility or packing differences—can dominate flexibility classification. Cluster-wise analysis reinforced the importance of structural context: prediction was most accurate for clusters 2 and 3, aligning with the hypothesis that flexibility prediction is facilitated in structural classes where RMSF variation is better captured by GI features.

In contrast, the regression task revealed a different optimal modeling strategy. While the attention-based CNN remained competitive, the recurrent neural network (RNN) achieved the highest R^2^ (0.537), outperforming all other tested architectures. This reversal in performance ranking suggests that continuous flexibility prediction benefits more from modeling sequential dependencies across the full GI vector, rather than focusing selectively on a few high-importance positions. Regression also proved more challenging than classification, with a tendency to underestimate high RMSF values, reflecting reduced sensitivity to extreme flexibility.

Cluster-specific regression outcomes were heterogeneous. Clusters 3 and 4 again exhibited the highest correlations between predicted and observed RMSF values, whereas cluster 1 showed no meaningful relationship—a result likely driven by both its α-helical enrichment and its small sample size (n = 19). This underlines the importance of considering both structural composition and data availability when evaluating model performance.

Together, these findings highlight that (i) GI descriptors encode structural and dynamical properties relevant to flexibility, (ii) model choice should be tailored to the prediction objective, with attention mechanisms favoring classification and sequence-aware architectures such as RNNs benefiting regression, and (iii) secondary structure composition exerts a strong influence on predictive accuracy. The combination of exploratory structural analysis and predictive modeling not only clarifies the strengths and limitations of GI-based approaches but also provides a roadmap for adapting architectures to specific tasks and structural contexts. Future work could expand on this framework by integrating additional sequence-derived or energetic descriptors, rebalancing underrepresented clusters, and exploring hybrid architectures that combine attention and recurrent processing for improved flexibility prediction across both categorical and continuous outcomes.

## Acknowledgement

This work was supported by the National Institutes of Health under Grant Numbers R01GM120238 and R01GM122804. The work is also partially supported by a start-up grant from Albert Einstein College of Medicine. Computational support was provided by Albert Einstein College of Medicine High Performance Computing Center.

## Author Contributions

F.V. Z.S., S.Y. and Y.W. designed research; F. V. and Y.W. performed research; F.V., and Y.W. analyzed data; F.V. and Y.W. wrote the paper.

---

**Supplementary Figure 1.**
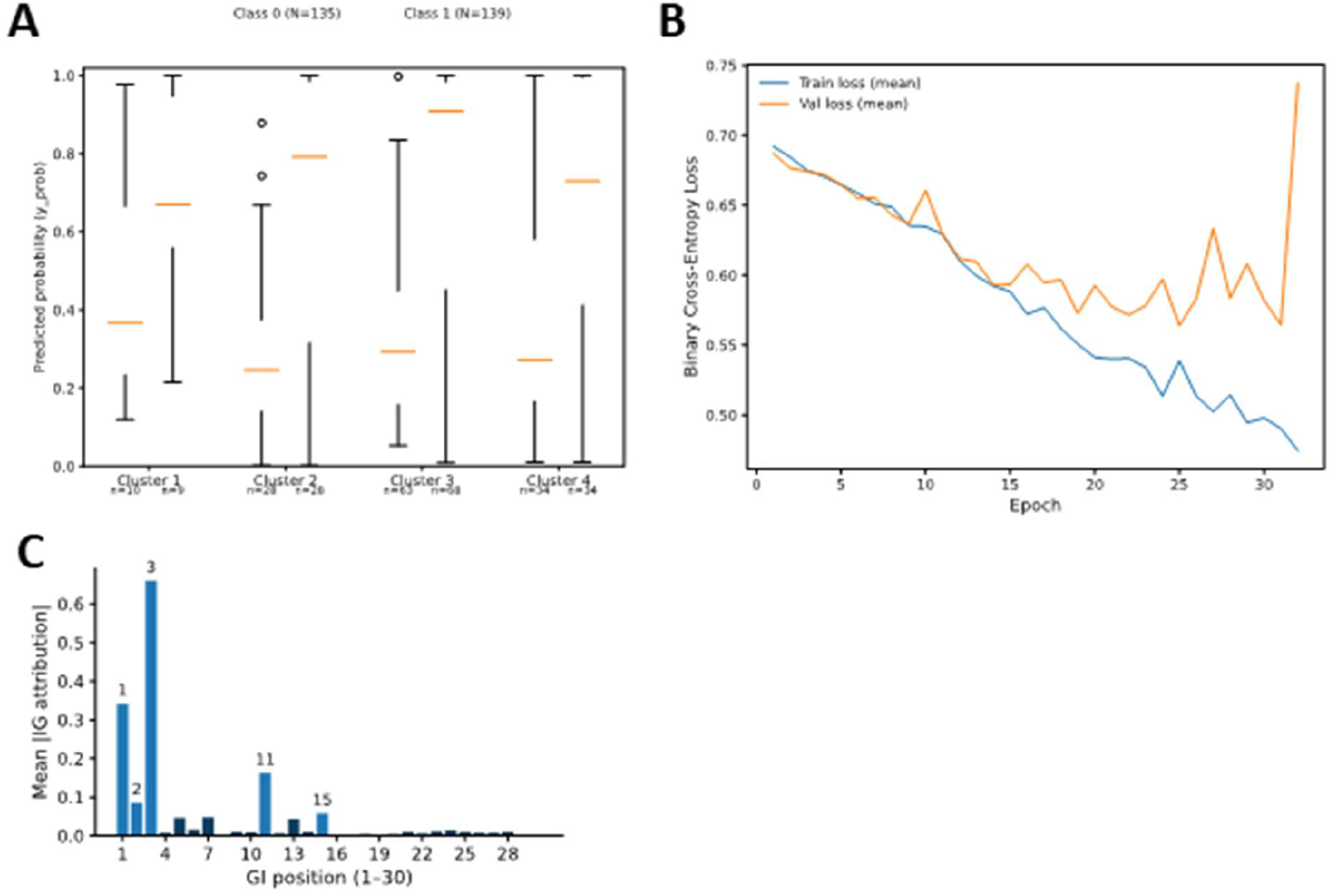
Additional analyses for the classification task. **A)** Predicted probability distributions across clusters for proteins labeled as non-flexible (class 0) and flexible (class 1). **B)** Training and validation binary cross-entropy loss curves across epochs. **C)** Feature attribution analysis using Integrated Gradients showing the relative contribution of each GI position (1–30) to the classification model.

**Supplementary Figure 2.**
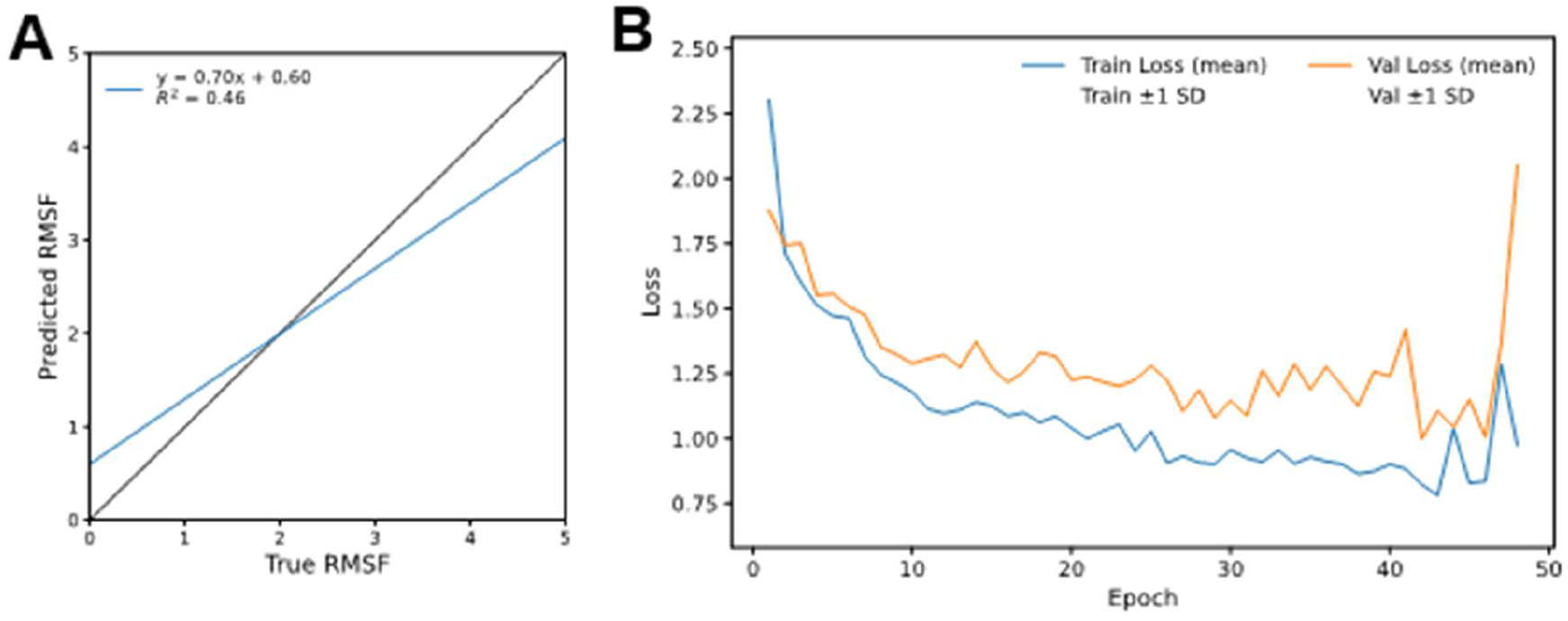
Additional analyses for the regression task. **A)** Zoomed scatter plot of predicted versus true RMSF values for proteins with RMSF ≤ 5Angstrom. **B)** Training and validation loss curves across epochs for the regression model, with shaded areas indicating standard deviation.

